# πDIA-CLIP: efficient identification of highly heterogeneous proteomics data via a generalized zero-shot framework

**DOI:** 10.64898/2026.02.09.704949

**Authors:** Yucheng Liao, Yongge Li, ZeXu Xiao, ChenChen Miao, Tao Yi, Xingpu Zhao, Yuanyuan Zhang, Han Wen, E Weinan, Cheng Chang, Weijie Zhang

## Abstract

Data-independent acquisition mass spectrometry has increasingly emerged as a cornerstone for characterizing highly heterogeneous biological systems, such as single-cell proteomics, metaproteomics, and spatial proteomics, offering unparalleled identification depth and quantification reproducibility. Current DIA analysis frameworks, however, require semi-supervised training within each run for peptide-spectrum match (PSM) re-scoring, which is prone to overfitting and lacks generalizability across diverse species and experimental conditions. Here, we present πDIA-CLIP, a generalized framework shifting the DIA analysis strategy from semi-supervised training to zero-shot cross-modal representation learning through integrating dual-encoder contrastive learning and encoder-decoder architectures to establish a unified, high-precision representation for spectral features and peptides. Notably, the generalized zero-shot nature of πDIA-CLIP facilitates an inference-only architecture, streamlining the analysis to achieve exceptional computational efficiency. Extensive evaluations across five distinct benchmarks demonstrate that πDIA-CLIP consistently outperforms existing tools, yielding an up to 44.6% increase in protein identification alongside a reduction in entrapment identifications reaching a maximal 52.5%. Furthermore, the enhanced identification depth facilitates the discovery of novel biomarkers and the elucidation of intricate cellular mechanisms.

## Introduction

Driven by the burgeoning demands of precision diagnostics and therapeutic development, proteomics has transitioned from a specialized technology to a high-throughput engine for biological discovery and clinical translation^1,2^. The ability to comprehensively characterize the proteome within highly heterogeneous biological contexts is now pivotal for identifying diagnostic biomarkers, monitoring treatment responses, and elucidating complex biological mechanisms^3–6^. To meet these rigorous requirements for depth and scalability, data-independent acquisition mass spectrometry (DIA-MS) has emerged as a cornerstone of high-throughput proteomics^7–10^, lauded for its high reproducibility^11^, sensitivity^12,13^, and high-throughput^14^. However, fully realizing the potential of DIA-MS is often hampered by the inherent complexity of its signal deconvolution, which remains a preeminent challenge in computational proteomics^15,16^. Unlike data-dependent acquisition (DDA) modes, DIA-MS employs isolation windows designed to encompass multiple m/z ranges, resulting in the concurrent fragmentation of multiple co-eluting precursor ions. This multiplexing gives rise to highly convoluted spectra where fragment-ion signals from disparate peptides are extensively interleaved, thereby confounding the definitive assignment of peptide-spectrum matches (PSMs) and limiting the depth of proteomic coverage^7,17^.

Over the past decade, MSFragger-DIA^18^, PECAN^19^, Spectronaut^20^, and MaxDIA^21^, have profoundly advanced the field of DIA proteomics, establishing the standard for reproducible proteome profiling. To analyze DIA-MS data, peptide-centric approaches rely on reference spectral libraries to query DIA-MS data with a standardized computational workflow: retention time (RT) calibration, extracted ion chromatogram (XIC) generation, peak-group scoring, PSM re-scoring, and false discovery rate (FDR) estimation. To distinguish true target peptides from decoy hits, earlier PSM re-scoring strategies relied on semi-supervised machine learning algorithms (e.g., Percolator^22^, mProphet^23^ or XGBoost^24^). These algorithms are predicated upon sophisticated feature engineering, which integrates diverse statistical metrics such as RT alignment, peak-shape correlation, and ionic intensity distributions to enhance identification confidence. However, the reliance on hand-crafted feature engineering inherently restricts the capacity to capture complex data nuances, thus compromising the sensitivity for identifying low-abundance proteins within complex background noise. Deep learning-based approaches, such as AlphaDIA^25^, DreamDIA^26^, DIA-NN^27^, Alpha-XIC^28^ and DIA-BERT^29^ leverage deep neural networks to extract high-dimensional features from raw DIA-MS signals without manual intervention, bolstering PSMs re-scoring performance and delivering substantially deeper proteome coverage.

Despite these advancements, existing frameworks still necessitate semi-supervised learning to repetitively optimize re-scoring models for each individual DIA-MS analysis run. By fundamentally assigning pseudo-labels to unannotated PSMs for iterative self-updating, this “run-specific” semi-supervised learning paradigm poses an inherent risk of overfitting. Specifically, biased initial annotations toward specific typologies drive the model into a detrimental, self-reinforcing loop, while uniform sampling strategies systematically marginalize rare species, inherently underfitting sparse, low-abundance targets. Beyond the risk of overfitting, the high computational cost required for repetitive optimization in semi-supervised learning framework severely bottlenecks high-throughput data analysis. Moreover, contemporary scoring metrics analyze peak groups in isolation, failing to capture the semantic associations between amino acid sequence and spectral data. These limitations highlight a critical need for a multi-modal framework capable of generalized PSM identification within sparse, high-dimensional space in a zero-shot manner, thereby circumventing the temporal burden of semi-supervised learning paradigm.

To circumvent these long-standing bottlenecks, we present πDIA-CLIP (<u>D</u>ata-Independent <u>A</u>cquisition with <u>Co</u>ntrastive <u>L</u>earning Integrated <u>P</u>roteomics), a generalized, end-to-end architecture for zero-shot PSM re-scoring, where “π” stands for the π-HuB project, a global initiative for proteomics research^30^. Diverging from traditional methodologies, πDIA-CLIP represents the first implementation of cross-modal contrastive learning in DIA-MS analysis, aligning discrete peptide sequences with corresponding chromatographic signal profiles. The core of πDIA-CLIP employs a hybrid architecture that synergizes a dual-encoder contrastive learning framework with a sophisticated encoder-decoder architecture. The dual-encoder component, which integrates transformer-based sequence encoder and a specialized spectral encoder, serves to align peptide sequences and XIC signals in a shared latent space. Concurrently, a decoder architecture leverages the aligned latent features to extract high-dimensional semantic abstractions. By harnessing pre-training on large-scale datasets, πDIA-CLIP effectively encodes the intricate, high-dimensional relationship between XIC profiles and corresponding peptides.

Furthermore, we extensively evaluated πDIA-CLIP across broad and heterogeneous DIA-MS proteomics cohorts, encompassing HeLa cell lysates, complex multi-species mixture, metaproteomic microbial communities, spatial breast cancer specimens and ultra-low-input single-cell preparations. Across these datasets, our evaluations demonstrate a marked expansion in detectable proteome depth and refinement of quantitative accuracy. Specifically, πDIA-CLIP achieved up to a 44.6% increase in protein identifications compared to existing software tools, while concurrently delivering a maximal 52.5% reduction in entrapment-based false identifications. Besides, driven by the zero-shot, inference-only architecture, πDIA-CLIP facilitates exceptional computational efficiency, achieving orders of magnitude speedup compared to existing deep learning-based methods. Beyond achieving superior benchmarks, πDIA-CLIP demonstrates exceptional translational utility in heterogeneous, practical scenarios, facilitating the discovery of novel biomarkers and the elucidation of intricate cellular mechanisms.

## Results

### Architecture and Analysis Workflow of πDIA-CLIP

πDIA-CLIP features a unified end-to-end architecture synergizing cross-modal contrastive learning framework^31^ with a dual-architectural paradigm comprising both dual-encoder and encoder-decoder architecture (Supplementary Information Fig. S1). The design rationale centers on the functional decoupling of cross-modal alignment and high-dimensional feature refinement. Specifically, the dual-encoder component utilizes a transformer-based sequence encoder and a specialized spectral encoder to project peptide sequences and XIC signals into a shared latent manifold for establishing foundational semantic correspondences. Complementing this global alignment, the encoder-decoder architecture serves as a discriminative engine, engineered to decode the intricate, non-linear dependencies between sequences and spectral signatures. This synergistic design couples the broad generalization of the dual-encoder with the high-resolution sensitivity of the encoder-decoder, facilitating precise PSM adjudication across heterogeneous datasets.

To circumvent the “run-specific” training constraints inherent in conventional DIA analysis, πDIA-CLIP employs a supervised contrastive learning pre-training strategy to internalize robust representations of peptide-spectrum correspondences (Fig. 1a). We curated a large-scale training dataset by processing DIA-MS data through DIA-NN, yielding an extensive collection of over 28 million high-confidence PSMs. This dataset encompasses four diverse species(*Homo sapiens*, *Mus musculus*, *Saccharomyces cerevisiae*, and *Escherichia coli*) and was acquired across five distinct MS platforms (e.g., Astral, TripleTOF; Supplementary Information Table 1). To bolster the discriminative performance of πDIA-CLIP against deceptive signals, entrapment PSMs are incorporated as negative sample during pre-training. This approach compels πDIA-CLIP to learn the subtle spectral nuances required to distinguish true targets from entrapments, ensuring accurate identification in a zero-shot manner. By pre-training on this large-scale dataset, πDIA-CLIP captures the intrinsic peptide-spectrum correspondences, fundamentally transcending the constraints of traditional feature engineering. Crucially, this entrapment-integrated pre-training strategy empowers πDIA-CLIP with zero-shot inference capabilities, enabling deep and accurate PSM identification with exceptional computational efficiency. To facilitate widespread academic and clinical adoption, πDIA-CLIP is deployed across a versatile ecosystem (Fig. 1a), supporting localized execution via a standalone workstation client, high-throughput processing through a scalable inference API, and automated down-stream interpretation integrated into LLM-driven agent analysts.

**Fig. 1:**
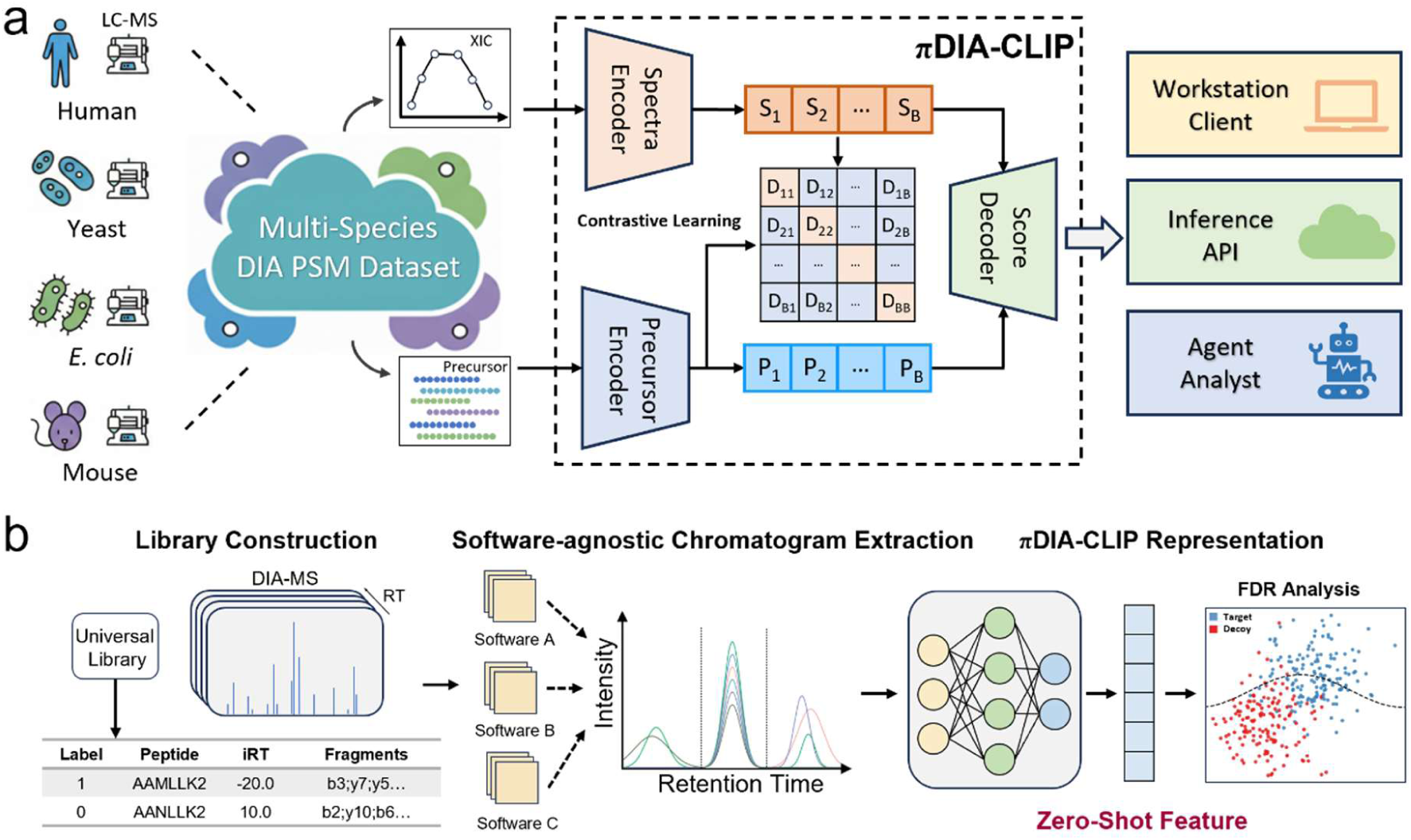
Overview of πDIA-CLIP Framework. a. Illustration of the supervised contrastive pre-training strategy. πDIA-CLIP leverages large-scale, diverse PSM dataset to learn robust and generalizable feature representations. By aligning precursors and XIC signals, πDIA-CLIP facilitates deep and precise peptide identification with zero-shot inference. b. Schematic representation of the πDIA-CLIP-integrated analysis pipeline. πDIA-CLIP is seamlessly incorporated into the standard DIA-MS workflow, specifically at the PSM re-scoring stage, to enable high-throughput and scalable proteomic profiling across diverse practical scenarios.

Leveraging the πDIA-CLIP, we established a comprehensive, peptide-centric workflow for end-to-end DIA-MS data analysis (Fig. 1b). This pipeline is designed to be software-agnostic, where RT calibration step can be performed by mainstream DIA identification tools. Subsequently, πDIA-CLIP extracts precursor and fragment XICs, performing PSM re-scoring and quantification within a zero-shot scenario. By imbuing the standardized workflow with extensive prior knowledge internalized during large-scale pre-training, πDIA-CLIP transcends biases inherent in individual experimental runs to ensure consistent, high-fidelity identification across varying experimental conditions.

### πDIA-CLIP enables high-sensitivity identification across diverse scenarios

To systematically evaluate the efficacy of πDIA-CLIP in proteome identification, we initially conducted rigorous benchmarking using HeLa cell lysates DIA-MS data across five distinct liquid-chromatographic (LC) gradients ranging from 0.5 to 4.0 hours (PRIDE ID: PXD005573; Fig. 2a)^32^. Crucially, to guarantee an unbiased assessment of zero-shot inference capability, all DIA-MS files used for validation were strictly excluded from the pre-training dataset.

**Figure 2.**
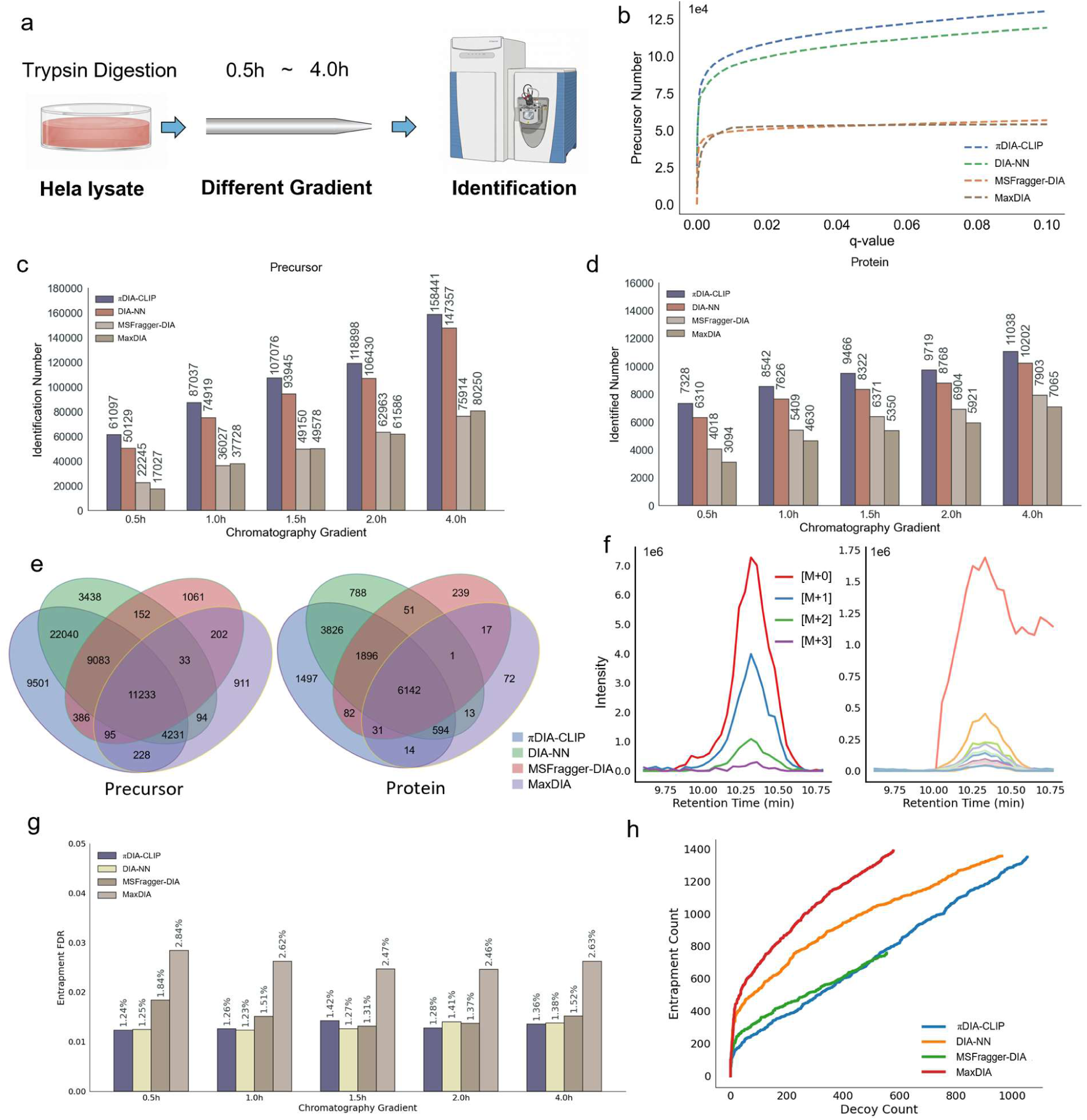
Comprehensive Benchmarking of πDIA-CLIP with HeLa Cell DIA-MS Data. a. Schematic of the benchmarking experimental design. b. FDR curve of precursor identification across various search engines at 90-minute LC gradient. c-d. Benchmarking results for peptide (c) and protein (d) identification across different LC gradient lengths. e. Venn diagram illustrating the overlap and unique counts of peptide and protein between πDIA-CLIP and other tools under 30-minute LC gradient. f. Representative XICs of target precursor that is uniquely identified by πDIA-CLIP. g. Evaluation of entrapment precursor FDR across different LC gradient lengths with decoy FDR < 0.01. h. Correlation between decoy and entrapment precursor counts at 120-minute LC gradient.

Evaluations of Hela cell lysate across LC gradients (Fig. 2c, d) revealed that the number of peptides and proteins identified by πDIA-CLIP markedly surpassed the benchmarks established by existing tools, such as DIA-NN, MaxDIA, and MSFragger-DIA. Taking the 90-minute LC gradient as a representative example, πDIA-CLIP achieved a 15.9% and 15.6% increase in precursor and protein identifications relative to DIA-NN, respectively (Fig. 2b). Visualization of target and decoy PSM score distributions further indicated that πDIA-CLIP constructs highly discriminative representations (Supplementary Information Fig. S2a). Consensus analysis of Venn diagrams further reveals a broad identification overlap between πDIA-CLIP and existing tools alongside the exclusive discovery of numerous high-quality unique peptides (Fig. 2e).

Furthermore, by leveraging narrow-window sampling and compressed chromatographic timescales, Orbitrap Astral MS instrumentation has fundamentally redefined the landscape of DIA proteomics, yielding substantially increased spectral density and complexity. As this unprecedented complexity poses a rigorous challenge for identification stability, we extended our evaluation to a highly multiplexed, multi-species consortium DIA-MS data acquired on this next-generation platform (PRIDE ID: PXD046444; Fig. 3a)^12^. This multi-species cohort consisted of proteomes from *H. sapiens*, *E. coli*, and *S. cerevisiae* at six distinct mixing ratios. Within this demanding acquisition environment, πDIA-CLIP identified the highest number of valid precursors (Fig. 3b), achieving at least 3.0% increase in precursor identifications and 6.7% increase in protein identification within 1.0% decoy FDR.

**Figure 3.**
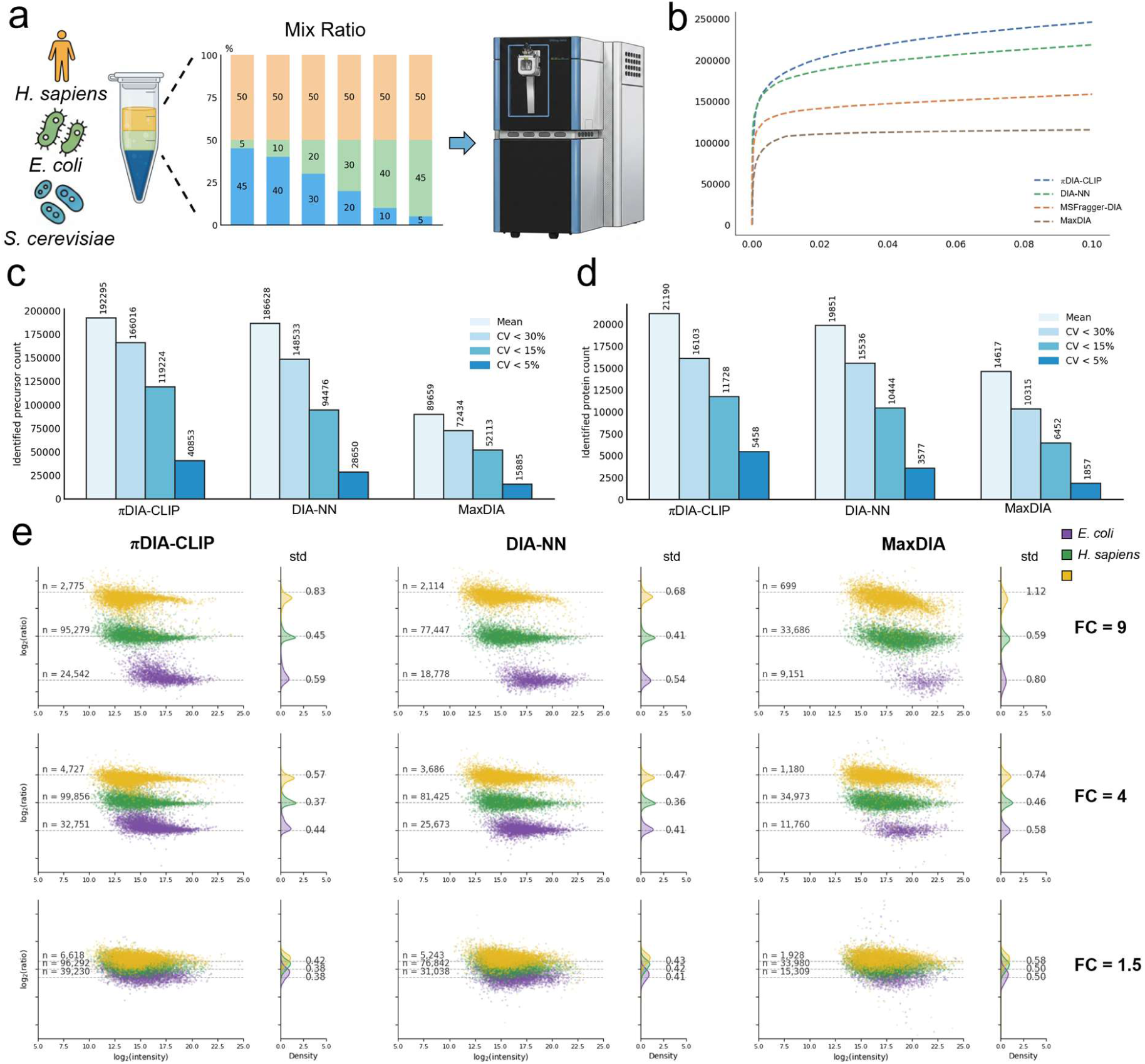
Benchmarking Results of Multi-species Samples. a. Experimental design of benchmarking dataset. The samples are generated by mixing known, varying ratios of cell lysates from *H. sapiens*, *E. coli*, and *S. cerevisiae*. b. FDR curve for precursor identification of various search engines with ratio = 10: 9: 1. c-d. Illustration of identification count of precursors (c) and proteins (d) within various CV on sample with ratio = 10: 9: 1. e-h. Comparison of relative quantitative precision across various tools. Scatter plots with marginal density distributions showing log-transformed ratios of quantified precursors. FC values correspond to theoretical mixing ratios of 50:45:5 to 50:5:45 (FC = 9), 50:40:10 to 50:10:40 (FC = 4), and 50:30:20 to 50:20:30 (FC = 1.5). Horizontal dashed lines demarcate the expected theoretical log2-transformed ratios.

Collectively, above results demonstrate that πDIA-CLIP overcomes the sensitivity bottlenecks inherent in heuristic feature engineering by leveraging the internalized, cross-modal representations to effectively resolve authentic signals. This approach establishes a robust computational foundation for high-throughput proteomics, facilitating deeper interrogation of the proteome across the latest generation of mass spectrometry platforms.

### Accuracy Assessment of πDIA-CLIP via Quantitative and Entrapment Performance

To assess the authenticity of precursor ions identified exclusively by πDIA-CLIP, we inspected XICs of representative precursor ions that remained undetected by established tools. For instance, in the 60-min HeLa cell lysates analysis, the +2 charged peptide ASDVHEVRK was uniquely identified by πDIA-CLIP, displaying exceptional peak symmetry and stringent fragment-ion co-elution profiles. This high signal fidelity was systematically corroborated by a global evaluation of XIC Pearson correlation coefficients. Precursors exclusively identified by πDIA-CLIP consistently exhibited higher average correlations across precursors, fragments, and cross-ion metrics (0.53, 0.30, and 0.13, respectively) compared to unique identifications of DIA-NN (0.49, 0.26, and 0.08), confirming the robust authenticity of these novel discoveries (Supplementary Information Fig. S2b). Additionally, this qualitative signal integrity is further corroborated by extensive additional XIC examples uniquely recovered by πDIA-CLIP across varying gradients (Supplementary Information Fig. S3).

Given the formidable challenge of accurate error control within DIA workflows and the failure of conventional search tools to maintain consistent FDR thresholds through entrapment procedures^33–35^, an entrapment experiment was conducted to rigorously validate the reliability and robustness of πDIA-CLIP. This methodology utilized a hybrid multi-species entrapment library comprising protein sequences from *H. sapiens*, *S. cerevisiae*, *E. coli*, and *M. musculus* to the analysis of HeLa cell lysates DIA-MS data (PRIDE ID: PXD005573; Fig. 2a), with any identified non-human peptides classified as entrapments. As illustrated in Fig. 2g, while maintaining a decoy FDR below 0.01, πDIA-CLIP yielded entrapment proportions comparable to existing tools across varying LC gradient lengths. Notably, application of more stringent decoy FDR thresholds revealed a marked reduction in entrapment FDR for πDIA-CLIP (Supplementary Information Fig. S2e, f). Specifically, under the 120-minute LC gradient at a highly conservative decoy FDR of 5e^-3^, πDIA-CLIP achieved 44.3% reduction in entrapment precursor counts compared to DIA-NN. In-depth investigation into the sensitivity toward decoy versus entrapment sequences further clarified this superior performance. Analysis of the 120-min gradient data revealed the highest sensitivity toward entrapment precursors in πDIA-CLIP, characterized by a consistently suppressed trajectory of falsely identified entrapment ions across the evaluated range and culminating in a maximal 52.5% reduction relative to DIA-NN (Fig. 2h).

Integrated quantitative evaluation across multiple datasets further substantiated the high accuracy of identified signals yielded by πDIA-CLIP. In the assessment of identification across varying coefficients of variation (CV) for precursors (Fig. 3c) and proteins (Fig. 3d) within the multi-species consortium, πDIA-CLIP maintained a substantial lead over existing software. Notably, in the high-precision regime (CV < 5%), πDIA-CLIP delivered 42.6% and 52.6% increase in precursor and protein identifications, respectively. Transitioning from intra-group reproducibility to relative quantitative fidelity, subsequent analyses evaluated the predefined mixing ratios inherent to this cohort. Prior to computing relative log2 ratios, rigorous data curation isolated robust precursors maintaining a coefficient of variation below 30%. Scatter plots mapping log2-transformed relative intensities against theoretical abundance illustrated exceptional concordance between πDIA-CLIP measurements and theoretical expectations (Fig. 3e). Crucially, the substantial gains in precursor identifications preserved quantitative fidelity, delivering a variance profile equivalent to established software tools. Most notably, evaluating mixtures with a 1.5-fold change (FC) reveals a distinctly tighter quantitative distribution for πDIA-CLIP, evidenced by narrower density profiles relative to existing tools. This systematic precision is further mirrored in the analysis of unique identifications in the HeLa dataset, where the quantitative results for precursors (Supplementary Information Fig. S2c) and proteins (Supplementary Information Fig. S2d) identified solely by πDIA-CLIP exhibited significantly tighter clustering alongside higher abundance levels relative to existing software.

These multi-dimensional evaluations collectively demonstrate the capacity of πDIA-CLIP for substantial increases in identification depth while maintaining high quantitative and statistical fidelity in zero-shot inference regimes. By effectively resolving the divergence between decoy and entrapment-based error estimates, πDIA-CLIP introduces a robust methodological advancement for accurate proteome interrogation, establishing itself as a foundational tool for high-fidelity, high-throughput proteomics across diverse DIA-MS platforms.

### Zero-Shot Inference Empowers Computational Efficiency

The advent of high-throughput MS, exemplified by the Orbitrap Astral, facilitates unprecedented data generation^12,36^. Despite the enhanced identification depth enabled by sophisticated deep learning architectures, the necessity for intensive training or optimization remains a substantial bottleneck. This lag between data acquisition and processing speed increasingly restricts the scalability of modern proteomic workflows.

πDIA-CLIP bridges this computational gap by shifting the DIA analysis strategy from repetitive semi-supervised training to a generalized zero-shot representation, offering a unified, inference-only solution for PSM re-scoring. The computational efficiency between πDIA-CLIP and existing deep learning tools was benchmarked using HeLa cell lysates analyzed with 0.5h chromatographic gradient (PRIDE ID: PXD005573). In this experiment, more than 26,000 MS/MS spectra were searched against a spectral library containing more than 4,000,000 precursors. To ensure hardware parity, all evaluations were conducted on a high-performance server equipped with 8 NVIDIA A100 (80GB) GPUs and dual Intel Xeon Platinum 8336C processors (2.30 GHz, 56 cores/112 threads). By obviating the computational burden of iterative optimization, πDIA-CLIP completed the entire workflow within 13 minutes, representing an 8-fold and a more than 100-fold acceleration in run time relative to AlphaDIA (1.7 h) and DIA-BERT (22.9 h), respectively (Supplementary Information Table 2). Furthermore, a granular assessment of PSM re-scoring and FDR estimation further underscored this efficiency, with πDIA-CLIP requiring only 2.8 minutes for these tasks. This performance contrasted sharply with AlphaDIA (31.7 min) and DIA-BERT (5.2 h) despite relying on high-performance GPU acceleration. This profound speedup stems from the zero-shot architecture of πDIA-CLIP, which requires only a single, highly efficient forward inference. This approach fundamentally bypasses the multiple computationally expensive epochs of model training and parameter updating mandated by existing models. Remarkably, πDIA-CLIP maintained its speed advantage even in CPU-only environments, with re-scoring and FDR stages (5 min) outperforming the GPU-dependent versions of existing frameworks. Such hardware-agnostic efficiency ensures superior platform extensibility, enabling high-performance proteomic analysis on diverse computational infrastructures independent of dedicated GPU requirements.

Collectively, these results establish πDIA-CLIP as an exceptionally efficient framework for high-throughput proteomics. By leveraging rapid, inference-only architecture, πDIA-CLIP provides a highly scalable and hardware-agnostic solution for modern proteomics, equipped to handle the massive analytical demands of large-scale MS datasets.

### Unlocking the Metaproteome via Zero-Shot Generalizability

Metaproteomics has emerged as a transformative tool for deciphering microbiome-host interactions and complex ecological networks by capturing community-wide functional profiles^37,38^. In practice, the wide-window acquisition of DIA-MS in metaproteomics yields highly chimeric spectra with overlapping fragment ions and composition-dependent interference, presenting a significant computational challenge for accurate profiling across diverse samples. Characterizing such immense taxonomic diversity within highly heterogeneous datasets necessitates robust generalizability, ensuring reliable identification of unseen taxa. To evaluate the generalizability of πDIA-CLIP across heterogeneous and previously unseen biological contexts, we benchmarked πDIA-CLIP utilizing a mixed metaproteomic cohort (PXD054415) comprising 32 species at equal cell counts^39^. The community spans immense taxonomic distances, including representatives from Archaea and Eukaryota alongside 25 bacterial strains and five bacteriophages. Since validated protein sequences for the Archaea, Eukaryota, and bacteriophages were unavailable in UniProt, library-free DIA-MS searching of our evaluation was restricted to a FASTA database of the 25 bacterial strains sourced from the original study^39^.

Comparative benchmarking on the PXD054415 cohort revealed superior identification generalizability for πDIA-CLIP at both precursor and protein levels, despite no prior exposure to these specific biological species. Total identifications of πDIA-CLIP reached approximately 11,000 precursors and 5,000 proteins, representing increases of at least 2.3% (Fig. 4a) and 2.0% (Fig. 4b) over existing tools, respectively. Species-wise analysis confirmed species-agnostic identification gains across the majority of microbial taxa (Fig. 4c). The improvements appeared particularly pronounced among low-abundance proteins, as evidenced by average 0.7% deviation in quantitative values and 13% reduction in standard deviation (Supplementary Information Fig. S4a). Furthermore, assessment of quantification stability revealed average 29.3% decrease in median CV values for πDIA-CLIP across all evaluated taxa, highlighting enhanced reproducibility in complex metaproteomic contexts (Supplementary Information Fig. S4b).

**Figure 4.**
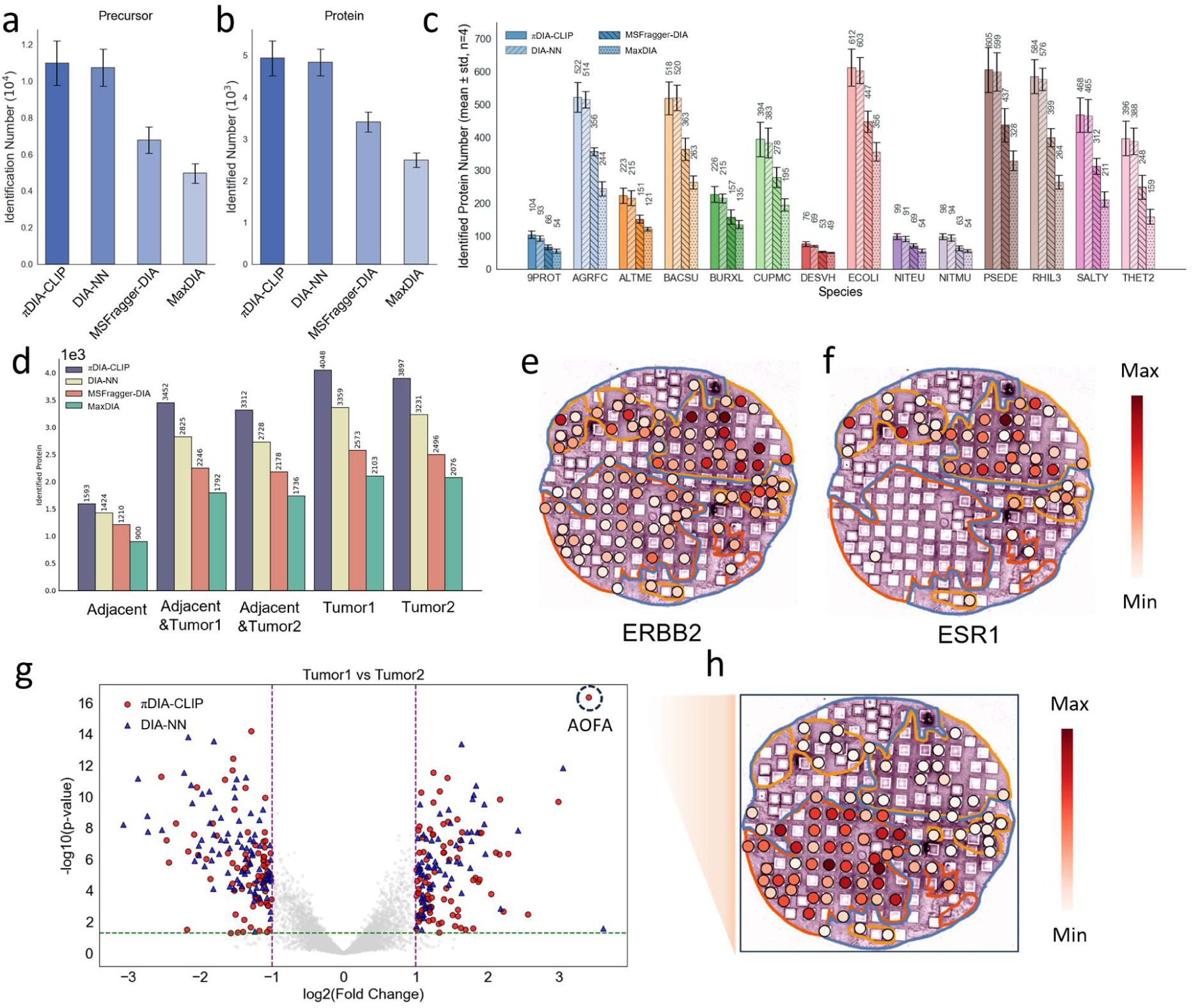
Benchmarking Performance in Metaproteomics and Enhanced Spatial Proteomics for Biomarker Discovery. a-b. Comparison of precursor (a) and protein (b) identification counts via different approaches on metaproteomic datasets. c. Species-wise identification counts across diverse microbial taxa. The distribution is derived from protein-level results presented in (b). d. Comparison of protein identification counts across different pathological regions. Only proteins identified in more than 50% of the tissue sections (LCM samples) are included in the statistics. e-f. Spatial distribution map of HER2/ERBB2 (e) and ER/ESR1 (f). Both are identified and quantified by the πDIA-CLIP. g. Volcano plots comparing differentially expressed proteins (DEPs) identified by πDIA-CLIP and DIA-NN between the two distinct breast cancer pathological regions. h. Spatial distribution map of the representative DEP (AOFA) uniquely identified by πDIA-CLIP between two distinct breast cancer pathological regions.

These results establish πDIA-CLIP as a robust and universally applicable framework for metaproteomic analysis across diverse, uncharacterized biological systems. The inherent generalizability of πDIA-CLIP eliminates the requirement for dataset-specific retraining, offering a scalable solution for large-scale proteomic investigations. By enabling the reliable identification of previously unseen taxa and peptide signatures, πDIA-CLIP facilitates the systematic exploration of the metaproteomic “dark proteome”, potentially unveiling novel functional drivers within complex ecological and host-associated networks.

### Deciphering Spatially Resolved Proteomes via πDIA-CLIP

Given the capacity to delineate protein profiles with precise spatial context, spatial proteomics has emerged as a transformative tool in various fields like tumor microenvironment research and clinical precision medicine^40–42^. To evaluate the practicality of πDIA-CLIP in these complex biological scenarios, a spatially resolved proteomics dataset generated by integrating laser capture microdissection (LCM) with liquid chromatography-mass spectrometry (LC-MS) is utilized to profile the proteomic landscape of breast cancer tissue sections (iProX ID: PXD045687)^43^. Pathological annotations derived from hematoxylin and eosin (H&E) stained images allowed for the precise resolution of spatial information across distinct pathological regions.

Comparative identification assessments across distinct pathological regions (Fig. 4d), demonstrated an average 19.3% increase in protein identification counts across all regions via πDIA-CLIP, providing a comprehensive molecular foundation for high-resolution spatial proteomic maps and biomarker discovery. To bolster data reliability, the analysis considered only proteins identified in at least 50% of tissue sections within a given pathological region. Subsequent analysis of key breast cancer classification markers, including epithelial growth factor receptor 2 (HER2/ERBB2) and estrogen receptor 1 (ER/ESR1)^44^, revealed pronounced spatial heterogeneity (Fig. 4e, f). The observed quantitative trends and spatial distributions aligned closely with the H&E-based pathological annotations, enabling the accurate stratification of two tumor subtypes: Tumor 1 (HER2+, ER-, PR-) and Tumor 2 (HER2+, ER+, PR+). These classifications showed high consistency with the findings of the original study^45,46^.

The enhanced identification depth provided by πDIA-CLIP facilitated the discovery of novel marker proteins through comparative analysis. Volcano plot (Fig. 4g) and Venn diagram (Supplementary Information Fig. S5a) analyses of differentially expressed proteins (DEPs) between two tumor regions yielded a 6% increase in the identification of significant markers. Subsequent gene ontology enrichment analysis of these DEPs revealed a highly significant enrichment of the epithelial-mesenchymal transition (EMT) pathway in Tumor 1, further corroborating its aggressive phenotype (Supplementary Information Fig. S5b). For instance, Amine oxidase [flavin-containing] A (AOFA), a DEP uniquely identified by πDIA-CLIP, displayed specific expression within Tumor 1 regions (Fig. 4h). Previous research associates AOFA with highly invasive phenotypes and the epithelial-mesenchymal transition process in breast cancer^47,48^, aligning with the aggressive pathological characteristics in Fig. 4b. Additional examples of DEPs uniquely recovered and spatially visualized by πDIA-CLIP are provided in the Supplementary Information Fig. S5c.

Collectively, these results demonstrate the robust applicability of πDIA-CLIP for the accurate interrogation of spatially resolved clinical proteomes. Beyond providing substantial gains in identification counts, the successful stratification of tumor subtypes and the discovery of novel, functionally pertinent protein markers underscore the capacity of deep biological discovery.

### Deep Profiling of Low-Abundance Proteins Resolves Single-Cell Heterogeneity

Single-cell MS-based proteomics is hindered by formidable technical bottlenecks, including ultra-low sample input, extreme signal sparsity, and pervasive data missingness^13,50–53^. To evaluate the performance of πDIA-CLIP in these low abundance samples, a dataset consisting of 12 technical replicates of HeLa single-cell samples was utilized (PXD049211)^54^. Furthermore, validation of πDIA-CLIP in practical biological contexts utilized a differentiation dataset encompassing 12 induced pluripotent stem cells (iPSCs) and 91 embryoid body (EB) cells (PXD049181)^54^. These datasets provided a rigorous basis for benchmarking identification depth, quantitative stability, and the resolution of biological heterogeneity at single-cell level.

Analysis of the HeLa single-cell samples demonstrated substantial superiority of πDIA-CLIP over other software tools, resulting in at least 33.6% and 44.6% improvements in precursor and protein identifications, respectively (Fig. 5a, b). An in-depth assessment of quantitative stability further demonstrated a substantial increase in the number of precursors and proteins identified by πDIA-CLIP across various CV thresholds, highlighted by approximately 3-fold and 4-fold improvement over DIA-NN within 20% CV, respectively (Supplementary Information Fig. S6b, c). XICs of representative precursors unique to πDIA-CLIP validated the expanded single-cell proteome (Fig. 5d). While single-cell signals suffered from peak tailing or intermittent signal gaps resulting from ultra-low ion counts (Fig. 5e, f), the accurate resolution of non-ideal signal profiles via πDIA-CLIP overcomes the sensitivity limits of conventional heuristic-based algorithms in high-noise environments, enabling a more coherent protein detection profile (Fig. 5c). Specifically, the bottom quartile (25%) of proteins identified by πDIA-CLIP exhibited 35.4% lower mean intensity compared to DIA-NN, while simultaneously achieving an 81.7% increase in identification count within this low-abundance regime.

**Figure 5.**
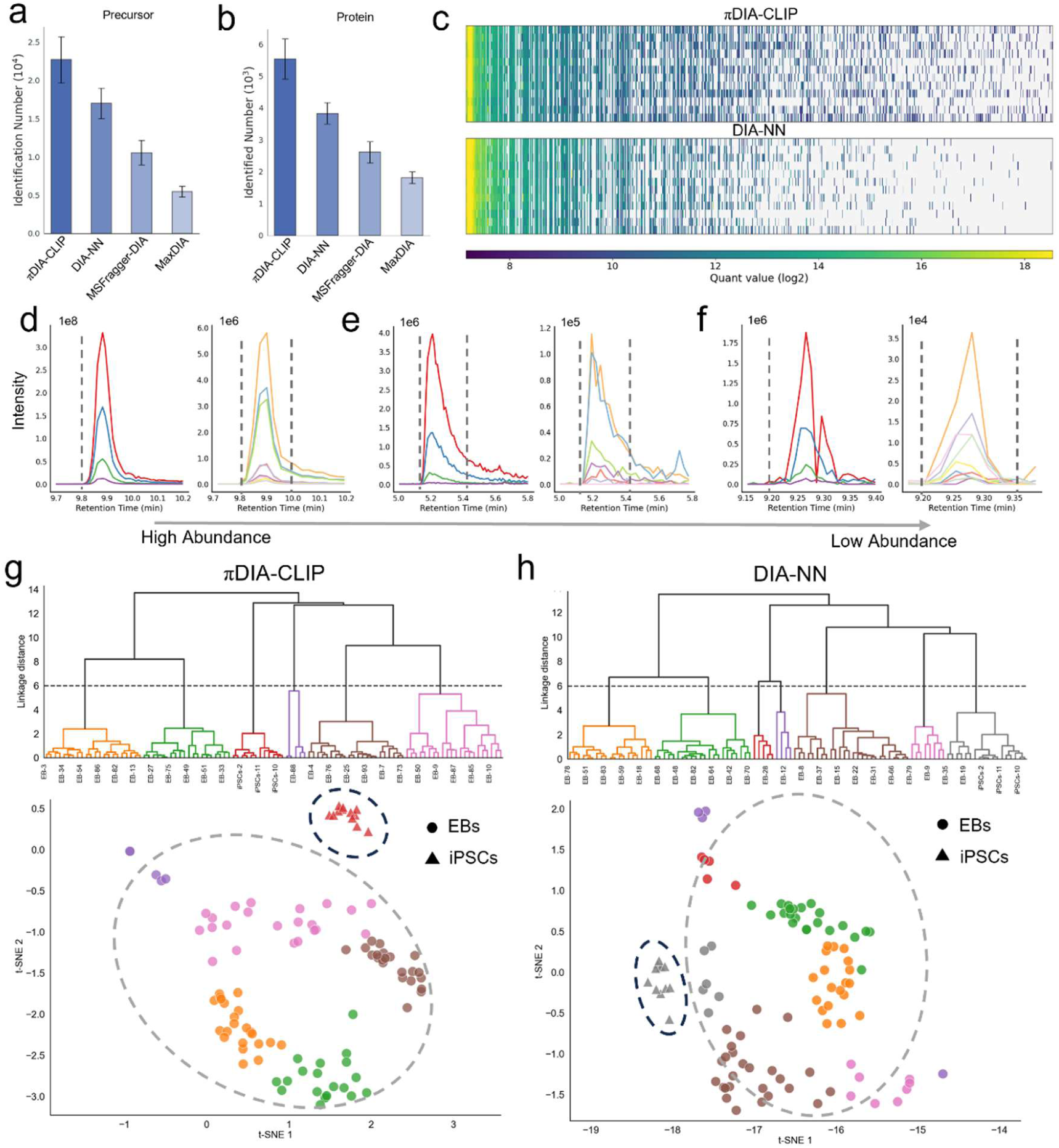
Benchmarking Evaluation and Biological Application of πDIA-CLIP in Single Cell Proteomics. a-b. Comparison of precursor (a) and protein (b) identification counts using Hela single-cell samples. c. Heatmaps of log2-transformed protein quantification values across samples using πDIA-CLIP and DIA-NN. Protein abundance is ranked from high to low (left to right). d-f. Representative XICs of target precursors uniquely identified by πDIA-CLIP, illustrating detection performance from high (d) to low (f) abundance. g-h. Unsupervised hierarchical clustering dendrograms and t-SNE dimensionality reduction plots of EBs and iPSCs. Dashed ellipses highlight the separation between EBs and iPSCs, comparing the biological resolution and sample classification accuracy of πDIA-CLIP (g) and DIA-NN (h).

Evaluation of the EB differentiation dataset demonstrates the biological utility of πDIA-CLIP. Following standardized preprocessing via zero-filling and intensity normalization, πDIA-CLIP yielded an expression matrix of 17,533 proteins across 103 cells, representing a 68% expansion in proteomic depth relative to the 10,432 proteins recovered by DIA-NN. Independent component analysis (ICA) applied to these quantification results produced low-dimensional embeddings for direct comparative assessment. These low-dimensional embeddings facilitated a crisp separation of iPSC and EB populations congruent with known lineage biology, whereas DIA-NN exhibited insufficient resolution for distinguishing these cell types (Fig. 5g, h). Quantitative assessment of cluster stability through hierarchical clustering and t-SNE^55^ visualization further substantiated the robustness of the πDIA-CLIP data. Protein expression heatmaps (Supplementary Information Fig. S6a) derived from these clusters further validated the accuracy of cell-type identities, highlighting the capacity of πDIA-CLIP to resolve heterogeneous scenarios.

Collectively, the superior identification depth and enhanced quantification stability achieved by πDIA-CLIP establish a robust computational foundation for high-fidelity single-cell proteomics. Improved resolution of non-ideal signal profiles and expanded coverage of the low-abundance proteome facilitate the elucidation of subtle biological heterogeneity within signal-sparse experimental landscapes.

## Discussion

This study introduces πDIA-CLIP, a cross-modal framework establishing novel identification approach in data-independent acquisition proteomics by transitioning from semi-supervised training models to a unified, generalized zero-shot inference architecture. By obviating the need for repetitive optimization, this approach eliminates the inherent risk of overfitting within constrained local datasets, while the inference-only architecture facilitates exceptional computational efficiency across heterogeneous biological scenarios, including metaproteomics, spatial proteomics, and single-cell proteomics. Mechanistically, πDIA-CLIP involves the first implementation of cross-modal contrastive learning within DIA-MS, where a dual-encoder framework aligns peptide sequences and multi-dimensional XIC signals within a shared latent space. Ablation studies further underscore the critical role of contrastive learning in fostering the generalizability of πDIA-CLIP, while simultaneously revealing substantial scope for architectural refinement to further enhance identification depth and sensitivity (see Supplementary Information Notes). Furthermore, by leveraging global prior knowledge from millions of PSMs, πDIA-CLIP ensures precise zero-shot identification and exceptional robustness across diverse instrumental platforms and biological scenarios, representing a foundational step toward truly scalable and generalizable proteomics informatics.

Systematic benchmarking across heterogeneous cohorts, including complex multi-species consortium, metaproteomic microbial communities, clinical breast cancer specimens and ultra-low-input single-cell preparations, demonstrates a marked expansion in both the depth and accuracy of the detectable proteome. Architecturally, πDIA-CLIP permits integration with various peptide-centric search engines, including MaxDIA, Spectronaut, and DIA-NN, for the purpose of PSM re-scoring, further enhancing the utility for generalized re-scoring across diverse computational workflows (see Supplementary Information Notes). Besides, the inference-only nature of πDIA-CLIP allows for flexible deployment across both CPU and GPU hardware, facilitating rapid integration into existing computational pipelines without the heavy computational overhead of training or finetuning.

Although the conventional FDR threshold serves as a golden standard, it fails to represent an absolute demarcation between authentic and spurious identifications. Due to inherent statistical fluctuations, there remain some authentic target signals in proximity to this threshold. Integrating expert experience to validate identifications within this critical boundary provides a “human in loop” framework for further elevating the performance ceiling of πDIA-CLIP. Establishing a closed-loop system where manually verified high-confidence identifications are fed back into a reinforcement learning framework is capable of facilitating the continuous refinement of the re-scoring model. This adaptive framework could transcend conservative FDR constraints to recover authentic signals, effectively pushing the sensitivity limits of πDIA-CLIP and enabling the discovery of novel biological features in deep-proteome profiles. Concurrently, a fundamental trade-off characterizes training data composition, as omitting entrapment sequences from the pretraining phase maximizes inference-level identification yields while inadvertently compromising the capacity to rigorously filter entrapment artifacts. Furthermore, expanding the training repertoire to encompass diverse post-translational modifications, non-tryptic peptides, top-down ultraviolet photodissociation (UVPD) fragmentations^56,57^ and ion mobility spectrometry (IMS) dimensions will be essential to enhance the versatility of πDIA-CLIP across complex, non-standard experimental landscapes.

## Methods

### Mass Spectrometry Files Used in Pre-training

Raw MS data utilized for training πDIA-CLIP were retrieved from the PRIDE database. Detailed information of these MS files is provided in Supplementary Information Table 1.

### Preparation of the Training Dataset

To accommodate the systematic discrepancies in XIC extraction and identification logic across software generations (Supplementary Information Fig. S7a, b), two distinct datasets were generated using different versions of DIA-NN (v1.7 and v2.0). The library-based identification procedure was performed by DIA-NN with parameters “--qvalue 0.01 --matrices --unimod4 --rt-profiling” (version 1.7) and “--matrices --gen-spec-lib --unimod4 --reanalyse --rt-profiling --report-decoys --out-lib-qvalue 0.01 --qvalue 0.01” (version 2.0). Prediction of the spectral library utilized a separate execution of DIA-NN (version 1.8.1) with parameters “--qvalue 0.01 --matrices --gen-spec-lib --predictor --fasta-search --min-fr-mz 200 --max-fr-mz 1800 --met-excision --cut K*,R* --missed-cleavages 1 --min-pep-len 7 --max-pep-len 30 -- min-pr-mz 300 --max-pr-mz 1800 --min-pr-charge 1 --max-pr-charge 4 --unimod4 --reanalyse --relaxed-prot-inf --smart-profiling --peak-center --no-ifs-removal”. Target precursors identified by DIA-NN served as positive samples, with all decoy precursors functioning as initial negative samples. To expand the negative sample space and enhance model robustness, an entrapment identification procedure was employed. All entrapment precursor identified by DIA-NN were incorporated as additional negative samples.

Extraction of XIC data utilized calibrated retention times and theoretical m/z value of both precursor ions and fragment ions, derived from DIA-NN. Precursor ion XICs originated from MS1 spectra, encompassing the monoisotopic precursor m/z and its three associated isotopic peaks. Fragment ion XICs were extracted from MS/MS spectra based on fragment m/z values. Peak groups searching employed a fixed m/z tolerance window of [ion m/z − 30 ppm, ion m/z + 30 ppm]. Within this window, only the peak with the m/z value closest to the theoretical expectation was selected. In instances where no peak was detected within the defined tolerance, the intensity was set to zero and the m/z error was recorded as 30 ppm. Following XIC extraction, both precursor and fragment XICs underwent intensity normalization and interpolation to a fixed dimension of 12 data points to standardize peak shape representation. Finally, the resulting peak groups were processed using a Gaussian smoothing algorithm to minimize signal noise.

### Re-scoring and Quantification Procedure of πDIA-CLIP

As illustrated in Figure S1, πDIA-CLIP is composed of three primary functional components: initial encoding phase, contrastive learning alignment stage, and specialized decoder for PSM re-scoring.

The initial encoding phase utilizes a dual-encoder architecture. Within a training batch of size B, precursor ions undergo transformation into feature representations P_1_, P_2_, …, P_B_. Besides, to handle the intrinsic relationship between precursor XICs and fragment XICs, πDIA-CLIP employs dedicated XIC encoding layers to separately process precursor XICs, fragment XICs, and concatenated XICs. These resulting features are then combined and processed to a representation of spectrum features S_1_, S_2_, …, S_B_.

The secondary stage involves a contrastive learning framework designed to align precursor and spectrum features within a unified, shared latent space. Optimization of πDIA-CLIP focuses on maximizing the similarity between (P_i_, S_i_) with label y_i_ = 1 (positive pairs) while simultaneously minimizing the similarity of (P_j_, S_j_) with label y_j_ = 0 (negative pairs).

Derivation of the PSM score utilizes a specialized decoder architecture. Encoded precursor and spectrum features are fed into a transformer-based decoder to generate a [B, dim_model] feature. In parallel, physical co-elution characteristics are explicitly incorporated through the calculation of Pearson Correlation Coefficient (PCC) matrix, subsequently projected into a [B, dim_model] feature via a multi-layer perceptron (MLP). Fusion of these two features occurs through element-wise summation, followed by a final MLP with a sigmoid activation function. This process yields a calibrated PSM score ranging from 0 to 1.

Calculation of quantitative abundance for each identified precursor relies on the arithmetic mean of Top-K areas of fragment XICs. While K conventionally equals 6, πDIA-CLIP adaptively selects the top-6 or all available fragment ions with non-zero areas, ensuring the inclusion of only informative signals in the final quantitative result.

### Loss Function of πDIA-CLIP

The training loss of πDIA-CLIP is a composite objective function designed to ensure both high-fidelity feature alignment and robust statistical discrimination. This loss function consists of a target-differentiated contrastive cross-entropy loss, a PSM classification loss, and a similarity margin loss, i.e.

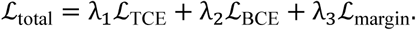

The target-differentiated contrastive cross-entropy loss (*L*_TCE_) imposes asymmetrical optimization goals on target and decoy/entrapment samples. Specifically, the loss function transforms the similarity matrix *S* into a modified matrix *S*′ by negating the rows associated with negative PSMs. Then row-wise application of traditional cross-entropy loss to the modified matrix follows the form of

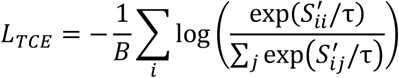

where τ serves as a temperature scaling parameter for matrix *S*′. Refinement of the final scoring output utilizes a PSM classification loss based on Binary Cross-Entropy (BCE), i.e.,

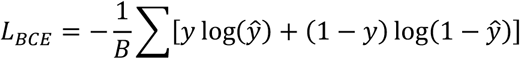

where *y* represents the ground-truth PSM labels and *ŷ* indicates the predicted PSM score vector. To ensure clear global separation between positive and negative identifications, πDIA-CLIP incorporates a similarity margin loss (*L*_margin_). This hinge-loss variant enforces a predefined margin (*m*) between the average similarity of target pairs (*μ_pos_*) and negative pairs (*μ_neg_*), i.e.,

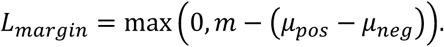

This component penalizes the model only upon insufficient statistical separation between these two populations, thereby fortifying the global discriminative capacity of the framework.

### Pre-training Parameters of πDIA-CLIP

πDIA-CLIP was jointly pre-trained using the AdamW optimizer with a weight decay coefficient of 1 × 10^-6^. To ensure stable convergence, a cosine annealing scheduler was implemented to dynamically modulate the learning rate, decaying from an initial value of 1 × 10^-4^ to a minimum of 1 × 10^-7^. The training spanned 40 epochs with a batch size of 4096. Dataset was partitioned into training and validation sets at a 9:1 ratio. Regarding the objective functions, we set the loss weights to λ_1_ = 1.0, λ_2_ = 1.0, λ_3_ = 5*e*^-3^, with the margin for similarity loss and the temperature scaling parameter setting to α = 0.5 and τ = 0.2 respectively.

### FDR Estimation

To evaluate the statistical confidence of PSMs, we employed the target-decoy strategy. In πDIA-CLIP, FDR control followed the Benjamini-Hochberg procedure^58,59^. Specifically, PSMs underwent initial ranking in ascending order of their scores to calculate raw FDR estimates. To ensure the monotonicity of FDR estimates, an adjustment was applied from the beginning of the ranked list, defined as:

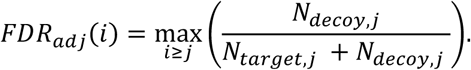

Target PSMs with an adjusted FDR below a predefined threshold (e.g., 0.01) were considered statistically significant identifications.

### Analysis of Spatial Proteomic Data

To evaluate the performance of πDIA-CLIP model in spatially resolved proteomics, we utilized an integrated computational workflow for the analysis of spatial DIA datasets, enabling the comparison of identification performance and molecular characterization across annotated tissue regions. Precursor-level identifications from πDIA-CLIP, DIA-NN, MSFragger-DIA and MaxDIA were mapped to a common spectral library to ensure standardized protein-level definitions, followed by protein quantification through precursor aggregation and outlier rejection. Spatial coordinates (x, y) were reconstructed for each tissue spot by matching sample metadata with position maps, and tissue-region annotations were verified by overlaying coordinates onto reference images.

Spatial protein distribution was visualized by normalizing protein abundance and overlaying these values on the tissue manifold. DEPs between tissue subregions (e.g., Tumor1 vs. Tumor2) were identified using Welch’s two-sample t-tests. Prior to statistical testing, quantitative outliers were removed to refine the abundance distributions. Significant DEPs were screened using comparative volcano plots, with FC > 2.0 and p < 0.01.

Functional interpretation of the spatial proteome was performed via Gene Ontology annotation using gseapy. To elucidate biological programs associated with specific tumor microenvironments, pathway enrichment analysis was conducted referencing the MSigDB Hallmark gene set collection. For these analyses, we employed threshold of FC > 2.0 and p < 0.01 to prioritize the identification of robust biological signatures. Enrichment significance was determined using Fisher’s exact test followed by FDR correction.

### Analysis of Single-cell DIA-MS Data

To standardize and analyze single-cell data across disparate search engines, we mapped identified precursors to a common human spectral library, followed by protein quantification through precursor-level averaging and outlier rejection. To account for technical variation in total signal, cell-specific protein abundances were normalized to a scale of one million units, and filtered to retain only proteins with normalized abundance exceeding 1.0 in at least two cells.

The resulting high-dimensional expression matrices were log-transformed and subjected to Independent Component Analysis (ICA) using the FastICA algorithm (random_state=3984) to identify the primary axes of cellular variation. Four independent components were extracted to serve as the basis for unsupervised hierarchical clustering via Ward’s method. The optimal clustering resolution was determined by assessing dendrogram linkage distances, with a threshold of 6 applied to partition cells into distinct subpopulations. To visualize these high-dimensional relationships in a lower-dimensional manifold, t-SNE algorithm (perplexity=75, random_state=254) was performed on the ICA component space.

### Benchmark setting

The benchmarking process included a comparative evaluation against several state-of-the-art software tools. Specifically, for the HeLa cell lysates and complex multi-species mixture, the evaluation was performed using DIA-NN (v2.0), MaxDIA (v2.6.7.0), and MSFragger-DIA (v4.1). Meanwhile, for the metaproteomics, spatial proteomics, and single-cell proteomics datasets, we utilized DIA-NN (v1.7.12), MaxDIA (v2.6.7.0) and MSFragger-DIA (v4.1). Reference proteomes for *H. sapiens*, *S. cerevisiae*, *E. coli*, and *M. musculus* were retrieved from the UniProt database.

For library-based search workflows, DIA-NN utilized spectral libraries predicted by DIA-NN (v1.8.1) based on the respective FASTA files, whereas MaxDIA employed its native library prediction engine. The predicted search space was constrained to peptide lengths ranging from 7 to 30 amino acids with precursor charge states between 1+ and 4+. Crucially, the MBR functionality remained disabled to ensure a standardized assessment of identification performance across all platforms. Analyses using MSFragger-DIA were conducted via the DIA-SpecLib-Quant workflow, with mass tolerances set to 10 ppm for precursors and 20 ppm for fragments. For all searches, carbamidomethylation of cysteine was considered as fixed modification. To ensure a standardized assessment of identification depth, protein-level identification was performed with q < 1.0, with a protein deemed identified upon at least one constituent peptide meeting the detection threshold.

Quantification assessments utilized precursor-level data across all evaluated platforms. Specifically, DIA-NN processing exclusively employed the direct quantification mode without MBR functionality. Precursor quantities were extracted from the ’Precursor.Quantity’ column in DIA-NN results and the ’Intensity’ column within evidence.txt files of MaxDIA. For protein-level quantification, a uniform approach involved calculating the arithmetic mean of constituent peptide intensities following the removal of outliers.

All other parameters for benchmarking software tools were maintained at their default settings, and comprehensive configuration files are provided in the Supporting Information.

## Data Availability

All raw data used for training and evaluation in this study were publicly available. All meticulously annotated training data and all results generated by πDIA-CLIP and benchmarking software were available at Zenodo (10.5281/zenodo.19059840).

## Code Availability

Code used for πDIA-CLIP inference is available at GitHub through the following link (https://github.com/Elcherneske/piDIA-CLIP). The executable software with a graphical user interface can be downloaded from https://github.com/Elcherneske/piDIA-CLIP/releases/. The agent system can be accessed through http://msrs1485008.bohrium.tech:50002/.

## Contributions

W.E., C.C. and W.Z. co-supervised the study. Y.Liao curated the training data, implemented the model, conducted the training and performed data evaluations. Y.Li provided expertise on the evaluation of single-cell data. Z.X. developed the graphical user interface for the software application. C.M. constructed the model-based agent system. T.Y., X.Z. and Y.Z. helped perform data evaluation. C.C. and W.Z. provided suggestions for bioinformatic analysis and data evaluations. H.W. and W.Z. provided computational resources. Y.Liao, C.C. and W.Z. drafted the manuscript. All authors reviewed and approved the final manuscript.

## Acknowledgements

This work was supported by the National Key R&D Program of China (2025YFA1309300) and National Natural Science Foundation of China (32088101). We acknowledge Zhifeng Gao from DP Technology for discussions about model training strategies. We thank Prof. Fangqing Zhao’s team for providing the spatial imaging information of breast cancer for evaluation.

## Ethics declarations

### Competing interests

The authors declare no competing interests.

## Notes

### Competing Interest Statement

The authors have declared no competing interest.

### Summary of Updates

main content updated; Figure 4 revised; author affiliations updated

